# Three-Dimensional Histology of Whole Zebrafish by Sub-Micron Synchrotron X-ray Micro-Tomography

**DOI:** 10.1101/392381

**Authors:** Yifu Ding, Daniel J. Vanselow, Maksim A. Yakovlev, Spencer R. Katz, Alex Y. Lin, Darin P. Clark, Phillip Vargas, Xuying Xin, Jean E. Copper, Victor A. Canfield, Khai C. Ang, Yuxin Wang, Xianghui Xiao, Francesco De Carlo, Damian B. van Rossum, Patrick La Rivière, Keith C. Cheng

## Abstract

Histological studies providing cellular insights into tissue architecture have been central to biological discovery and remain clinically invaluable today. Extending histology to three dimensions would be transformational for research and diagnostics. However, three-dimensional histology is impractical using current techniques. We have customized sample preparation, synchrotron X-ray tomographic parameters, and three-dimensional image analysis to allow for complete histological phenotyping using whole larval and juvenile zebrafish. The resulting digital zebrafish can be virtually sectioned and visualized in any plane. Whole-animal reconstructions at subcellular resolution also enable computational characterization of the zebrafish nervous system by region-specific detection of cell nuclei and quantitative assessment of individual phenotypic variation. Three-dimensional histological phenotyping has potential use in genetic and chemical screens, and in clinical and toxicological tissue diagnostics.

**One Sentence Summary:** Synchrotron X-ray micro-tomography can be used to rapidly create 3-dimensional images of fixed and stained specimens without sectioning, enabling computational histological phenotyping at cellular resolution.

## Introduction

Histology has been used for more than a century to visualize cellular composition and tissue architecture in millimeter- to centimeter-scale tissues from diverse multicellular organisms (*1*). Its diagnostic power is dependent on the detection and description of changes in cell and tissue architecture. Normal and abnormal cytological features indicative of physical, inflammatory, and neoplastic causes of disease are readily distinguished using histology (*2*).

Despite its power, histology has practical limitations that interfere with sample throughput in large-scale studies of genetic and chemical phenotypes. Chief among these is the requirement to physically section specimens. Physical sectioning introduces tissue loss and distortions that make complete reconstruction of tissues from serially sectioned histology challenging (*3*). Even with expert proficiency using histological tools and equipment, slice thickness is limited to ∼5 µm due the technical demands and physical constraints of cutting paraffin blocks. This leads to incomplete sampling and imperfect visualization of elongated structures such as vessels and large cells, which frequently extend beyond the 5 µm section thickness. As a result of these issues, only a small fraction of any given tissue sample is studied in histology. Moreover, it is intractable to generate complete sets of sections for large numbers of whole organisms (as needed for a genetic or chemical screen) and impractical to align them for volumetric analysis. A routine method for histological phenotyping that avoids these problems and enables comprehensive three-dimensional (3D) analysis of whole organisms would enable transformative large-scale studies.

Optical imaging modalities resolve 3D sub-micron features but depend on sample transparency and/or require physical slicing to maintain resolution without diffraction or loss of signal (*4*–*7*). Serial block-face scanning electron microscopy has been used to image and reconstruct a mouse neocortex (*8*) and the larval zebrafish brain at nanometer-scale (56.4 × 56.4 × 60 nm^3^ resolution from 16,000 sections) (*9*). These studies have made it possible to define precise neurological and circulatory relationships across the entire brain. However, this method is time-consuming and thus infeasible for interrogating replicate specimens required for toxicology or genetic screens. For example, a multi-beam scanning electron microscope, imaging a cubic mm of tissue at 20 nm^3^ resolution can require ∼3 months of continuous imaging (*10*).

X-ray micro-tomography (micro-CT) is a potential means of achieving complete, 3D histological phenotyping for large numbers of specimens. Micro-CT is commonly used to study hard, mineralized tissues like bone and fossils (*11, 12*). Soft-tissue imaging typically requires contrast-enhancing, heavy-metal stains such as osmium tetroxide, iodine, phosphotungstic acid (PTA), or gallocyanin-chromalum (*13*–*15*). Phase-based synchrotron imaging of unstained samples allows the study of development in live whole-embryos over time (*16*) but metal-based stains are required to achieve the contrast between subcellular components that is necessary for histology-like evaluation. Table-top micro-CT devices have been used to interrogate soft-tissue structure in diverse stained specimens (*17*–*19*), but these devices utilize polychromatic, low-flux X-ray tube sources that limit resolution and throughput. Synchrotron X-ray sources have monochromatic, high-flux beams (*20*) that allow rapid imaging of mm-scale samples such as insects, vertebrate embryos (*14, 15*), and mouse somatosensory cortex (*10*). For larger samples such as whole mice or whole mouse organs, imaging is performed at lower resolution (∼10 µm^3^) (*21, 22*), which does not permit cellular analysis. Conversely, nano-CT enables imaging at higher resolution (∼100 nm^3^) but does not provide sufficient field-of-view for whole mm-scale samples. To our knowledge, no existing method has the combination of throughput, resolution, field-of-view, and soft-tissue contrast necessary for whole-organism 3D phenotyping.

Here, we applied modifications of synchrotron X-ray micro-CT to create a virtual, 3D histological analysis of a whole model vertebrate organism, the zebrafish, at sub-micron resolution. We chose the zebrafish based on its relatively small size as a vertebrate organism (juveniles are <3 mm in diameter) and its wide range of tissues that facilitate evaluating metal-based stains for micro-CT based histological analysis. These customizations enable complete histological phenotyping of whole organisms to probe the relationships between genotype, environment, and morphology (*23*–*25*).

## Results

### Whole-Organism Synchrotron X-ray Histo-Tomography

The demanding aspects of high-throughput phenotyping on whole organisms led us to consider synchrotron based micro-CT. This is due to the combination of high flux and adjustable acquisition parameters that are typical of a synchrotron X-ray source. Fine-tuning of X-ray energy and bandwidth, along with optimizing image-acquisition geometry, allow optimization of the volumetric reconstructions of optically opaque zebrafish at isotropic resolutions (equal in all three dimensions). Of note, synchrotron X-ray flux is orders of magnitude more brilliant than commercial tube sources, greatly shortening imaging times (*20*).

Micro-CT studies were performed on the sector two bending magnet, B hutch (2-BM-B) of the Advanced Photon Source at Argonne National Laboratory for both single-energy and multi-energy acquisitions (**Figure 1**). We selected beam energies for monochromatic imaging studies to optimize contrast-to-noise ratio across a range of sample diameters and concentrations of tungsten (**Figure S1**). A beam energy of 13.8 keV was used for larval samples and 16.2 keV for juveniles due to larger thickness (see Methods). Sample-to-scintillator distance was adjusted to modify the magnitude of phase contrast-based edge enhancement (**Figure S2**). A sample-to-scintillator distance of 30 mm provided sufficient edge enhancement to achieve histology-like contrast. For whole-organism imaging, the zebrafish were vertically translated to capture segments over the full length of the sample. Reconstructions were combined into a full 3D volume (**Movie S1**).

**Figure 1.**
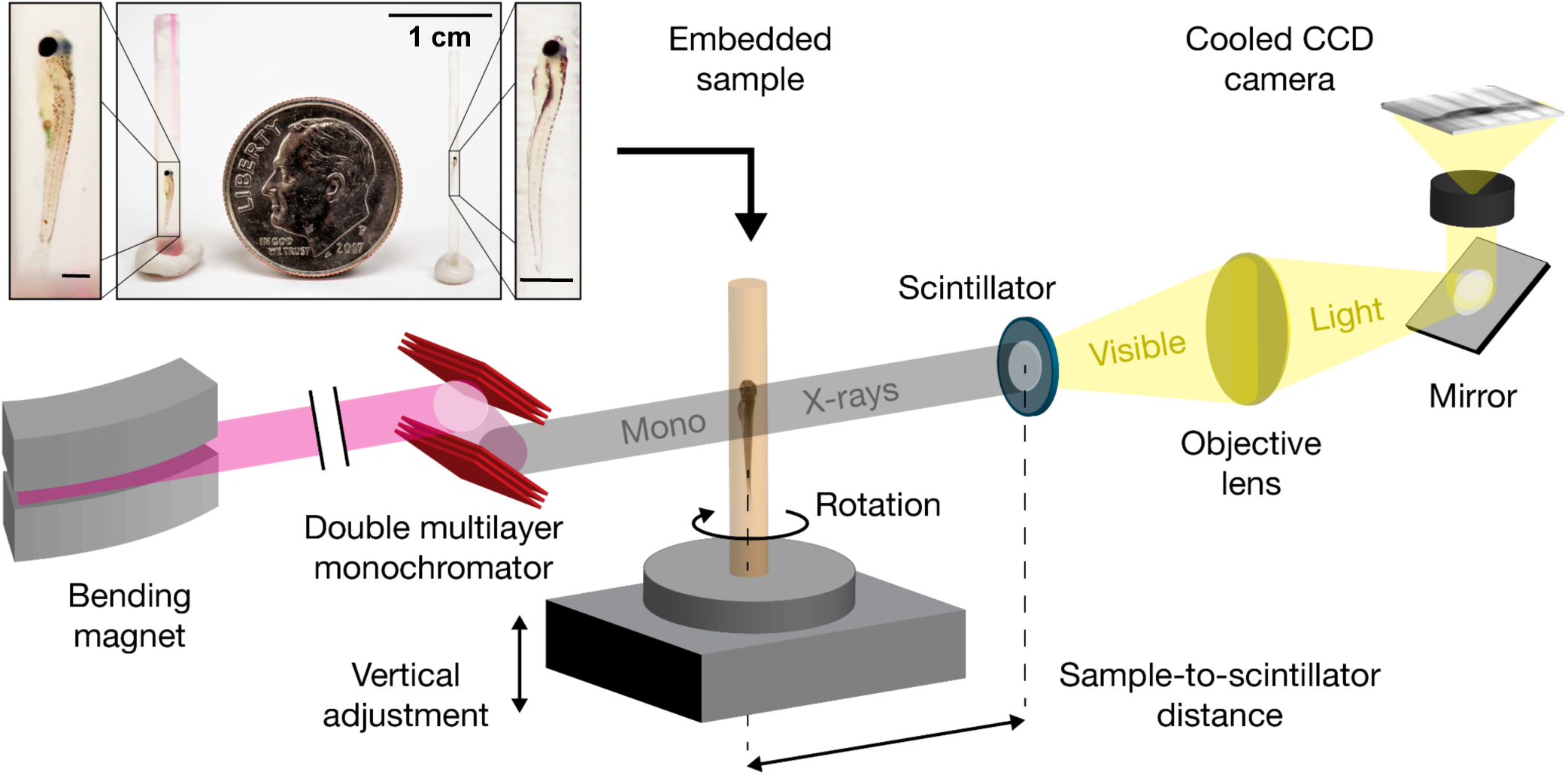
Schematic for synchrotron X-ray micro-tomography of whole zebrafish. Quasi-parallel X-rays from beamline 2-BM-B are used to acquire projection images of an intact, fixed, and PTA-stained whole zebrafish. Total imaging time is ∼20 minutes per 2048 slice-section for monochromatic acquisitions (sample-to-scintillator distance = 30 mm) and ∼20 seconds per 2048-slice section for pink-beam acquisitions (sample-to-scintillator distance = 25 mm). The top inset in **Figure 1** shows a juvenile and larval sample relative to a dime (diameter = 17.9 mm). For clarity, the polyimide tubing was removed prior to photography, but this not necessary for successful X-ray image acquisition. Scale bars in the specimen insets are 1 mm.

Large-scale phenotyping studies that focus, for example, on a variety of mutants or chemical exposures, require very short imaging times to facilitate throughput. Monochromatic acquisitions took ∼20 minutes, ∼36-fold faster than acquisition by commercial tabletop sources (e.g., ∼720 minutes for Xradia 500 series machines) (*15*). The use of polychromatic “pink” beam increases X-ray flux, which greatly reduces exposure times. Each pink-beam acquisition took ∼20 seconds, ∼60-fold faster than monochromatic acquisition and ∼2,000-fold faster than commercial sources. Monochromatic acquisitions have better image quality than pink-beam, as defined by signal-to-noise ratio and pixel intensity profile (**Figure S3**). Notably, no image degradation over years of repetitive synchrotron imaging was detected (*26*). This high degree of sample stability across multiple scans suggests the potential for pink-beam pre-screening followed by higher resolution monochromatic reacquisition.

### A Digital Zebrafish at Cell Resolution

Here we present a digital zebrafish that is stained in all tissues by PTA, at a sub-cellular voxel (volume pixel) resolution, with a field-of-view of multiple millimeters. The similarity of our digital images to conventional histological outputs is demonstrated by representative transverse (axial), sagittal, and coronal cross-sections for larval (4 days post-fertilization, dpf) and juvenile (33 dpf) zebrafish imaged at 0.743 and 1.43 µm^3^ isotropic voxel resolution, respectively (**Figure 2**). Volume renderings illustrate the orientation of individual planes of sections in 3D. Full sets of cross-sections of the zebrafish in all three orientations show how micro-CT data can be used to phenotype the full tissue volume to guide further analyses (see Movies S2-3).

**Figure 2.**
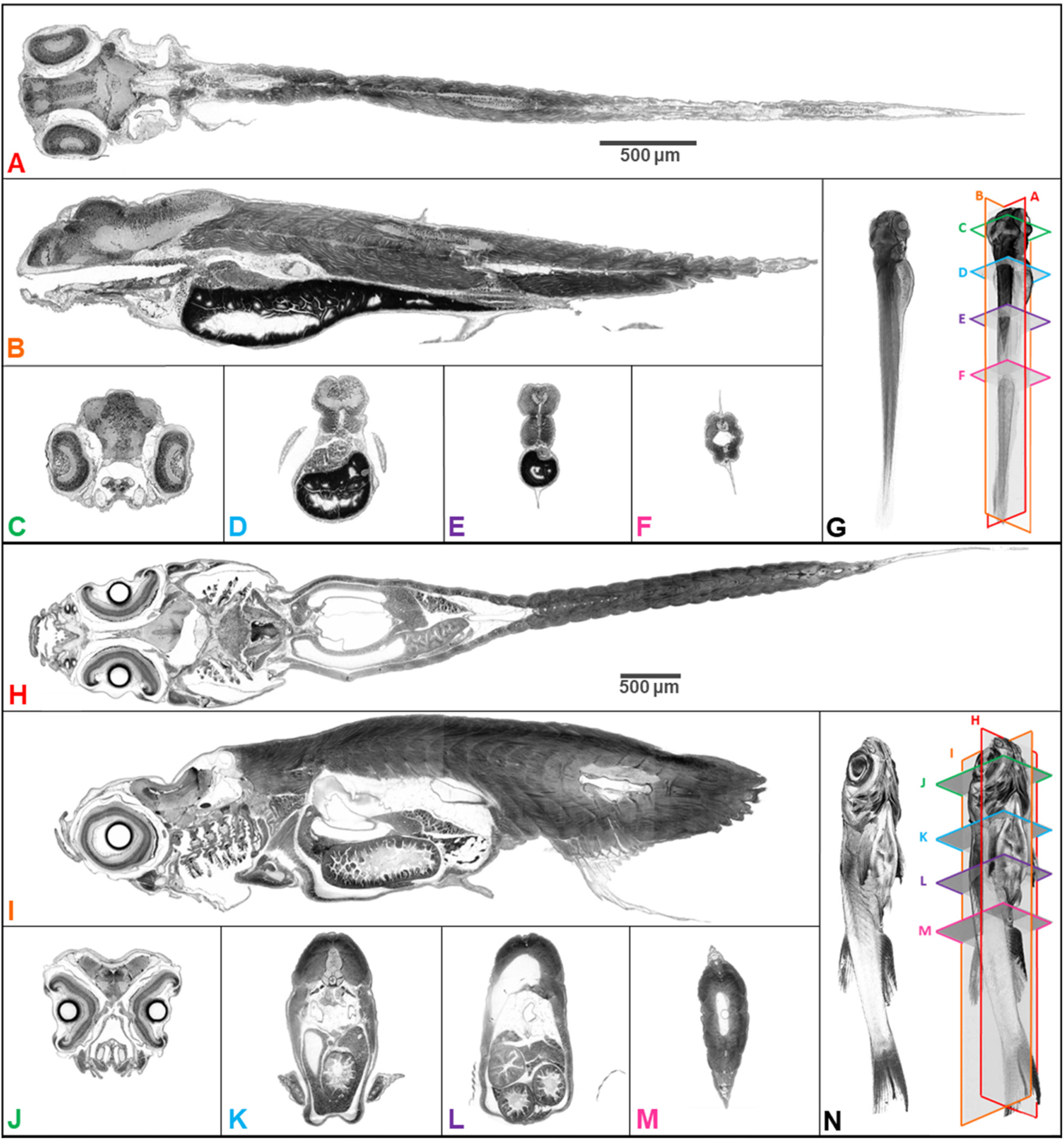
Whole organism imaging of PTA-stained zebrafish at cell resolution enables histology-like cross sections. Coronal (A, H), sagittal (B, I), and axial (C-F, J-M) cross sections of 4 dpf larval (A-G) and 33-dpf juvenile (H-N) wild-type zebrafish acquired using synchrotron X-ray micro-tomography at 0.743 μm^3^ and 1.43 μm^3^ isotropic voxel resolution, respectively. 3D volume renderings (G, N) show the cross-sections in relation to the whole organism. In contrast to histology, the cross sections are single voxel in thickness and can be obtained in any plane (including oblique cuts) after imaging. Complete cross-sections in the orthogonal directions for both fish are provided (**Movies S2 and S3**). Images are presented to match histological convention of dark cell nuclei (higher attenuation is darker).

A direct comparison between histo-tomography and histology can be achieved by creating ∼5 µm maximum intensity projections to mimic the thickness of conventional histology sections. These visualizations from ∼3.5 mm-long larval (5 dpf) and ∼1 cm-long juvenile (33 dpf) zebrafish with their corresponding hematoxylin and eosin (H&E) stained sections from age-matched zebrafish reveal a similar ability to recognize tissue and cell types (**Figure 3**). The resolution is comparable with that of optical histology using a 10X objective lens. The achieved resolution allows the recognition of individual nuclei in the larval brain (**Figure 3A-B**) and the measurement of regular spacing of striations (average = 2.16 μm, SD = 0.55 μm, n = 293 measures) in the skeletal muscle encircling the swim bladder (**Figure S4**), in accord with previous measurements (*27, 28*).

**Figure 3.**
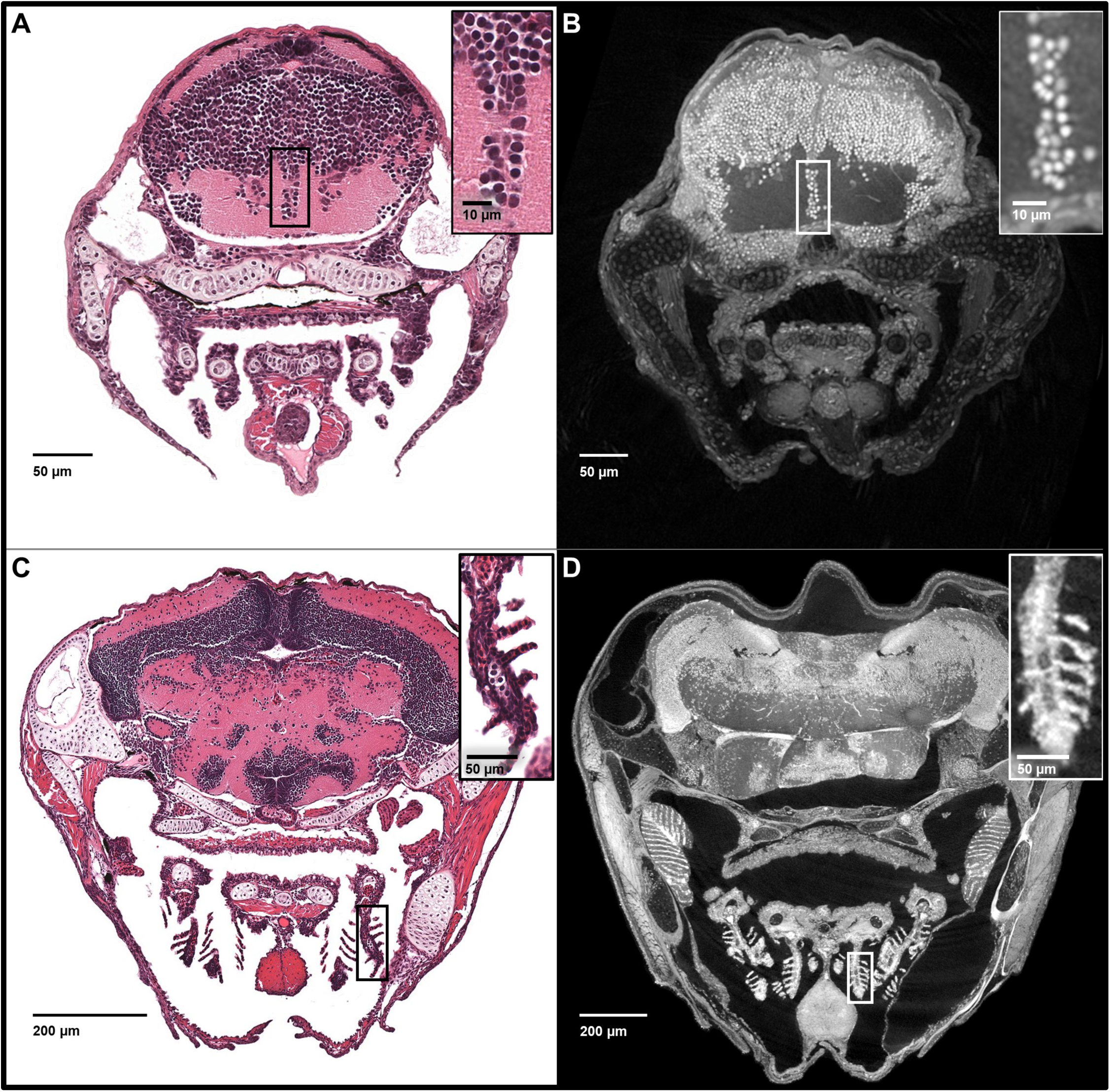
Synchrotron X-ray micro-tomography imaging produces sets of histology-like images. *Top:* Comparison of larval (5 dpf) zebrafish obtained from a 5 μm thick histology section and micro-tomography data (A and B, respectively). The micro-tomography data is 0.743 μm^3^ isotropic voxel resolution, and a maximum intensity projection (MIP) of 7 slices is shown to correspond with the thickness of histological sections. *Bottom:* Comparison of juvenile (33 dpf) zebrafish in both 5 μm histology section and microtomography (C and D, respectively). This data is 1.43 μm^3^ isotropic voxel resolution, and a MIP of 3 slices is shown. Insets show detail of both brain cell nuclei (A-B) and delicate gill structure (C-D). The images demonstrate the near histological resolution of X-ray histo-tomography, including the ability to distinguish individual cell nuclei. Although natural variation in size is observed in age matched fish (panel C length = 7.8 mm, panel D length = 10 mm) individual features are consistent.

The visualization of complex 3D tissue structure is made possible in the digital zebrafish. In histological images, limited aspects of the arrangement of gill cartilage, capillaries, and epithelial cells are visible, but the overall shape of gill lamellae is not apparent (**Figure 3C**). Reconstructions of juvenile zebrafish at 1.43 µm^3^ resolution similarly reveals the two-dimensional (2D) structure of gills (**Figure 3D**). While both traditional histology and histo-tomography have the resolution needed to distinguish cellular features in 2D slices, only the latter is able to reveal complex 3D tissue architecture. The use of micro-CT makes it possible to interrogate primary and secondary lamellae from multiple angles without spatial distortion, revealing the delicate leaf-like structure of gill filaments on any pharyngeal arch (**Movie S4**). The whole animal reconstructions provide a level of organismal and anatomical context that is not practical using histology alone.

### 3D Histological Analysis of Zebrafish Microanatomy

Conventional histological analysis requires time-consuming physical slicing through samples in one plane. Complete sectioning of a 1 mm^3^ tissue sample embedded in plastic would require hundreds of physical slices. The throughput of histological sectioning of zebrafish samples is especially slow since the block has to be soaked in ice water for ∼10 to 15 minutes between sets of 10-12 section ribbons (*29*). Thus, full volume analysis of whole zebrafish using histology is prohibitively slow.

The isotropic nature of our data enables reslicing in any plane of section after imaging. This is particularly useful for interrogating convoluted structures such as the gut. Dynamic reslicing (cutting the same volume digitally multiple times after image acquisition) allows the study of either longitudinal or perpendicular virtual sections of the gut at cellular resolution (**Figure 4**). In the histological phenotyping workflow, this facilitates complete evaluation of convoluted structures, which is intractable with conventional approaches.

**Figure 4.**
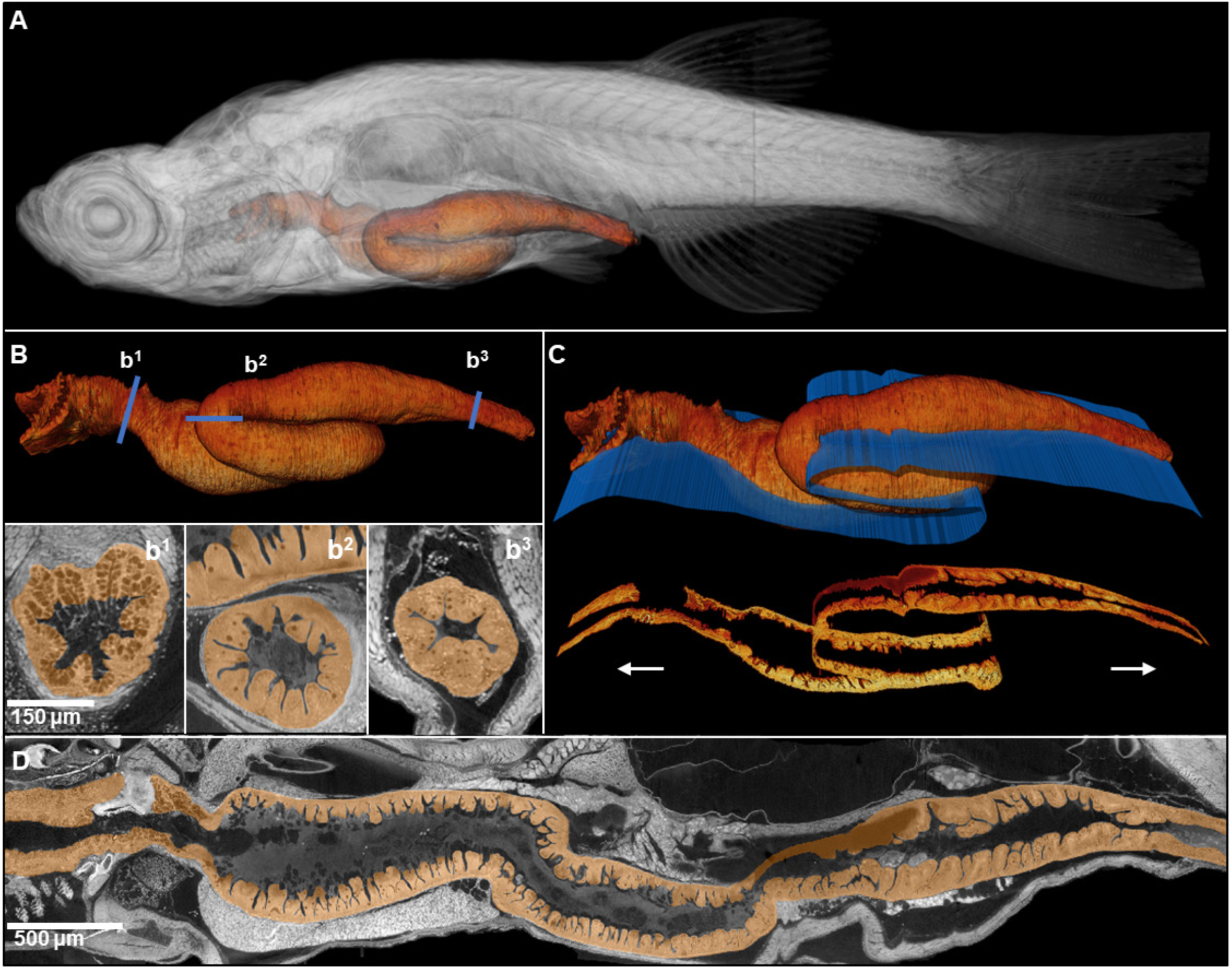
Comprehensive histological cross-sectioning of convoluted structures in juvenile zebrafish. A 3D rendering of a whole juvenile (33 dpf) zebrafish is presented with highlighted gastrointestinal (GI) tract (A). The GI tract exemplifies a convoluted structure for that can be unraveled for histological analysis. The isolated GI tract is displayed (B). Isotropic resolution of data permits virtual slicing at any angle (B1-B3) without a decrease in resolution, allowing comprehensive histology-like visualization despite its tortuous nature. A spline-based reslicing method for visualizing serpentine organs across their entire path is also shown (C). A nonlinear cutting plane (blue) follows the structure of interest, allowing the cut to render a structure’s total length onto a single plane (D).

Virtual sectioning and 3D rendering can be used to study any tissue in the fish, including neuronal cells in the eye, cartilaginous rudiments of the notochord, the squamous patch of the dorsum of the pharynx, nucleated red blood cells in the heart, and the pneumatic duct and goblet cells of the intestine of whole larval (5 dpf) and juvenile (33 dpf) specimens (**Figure S5** and **Figure 5**, respectively). In sum, the digital zebrafish allow tissues and cell nuclei to be visualized across organ systems, such as the integumentary, hematopoietic, respiratory, genitourinary, musculoskeletal, gastrointestinal, cardiovascular, nervous and sensory systems (**Movie S5**).

**Figure 5.**
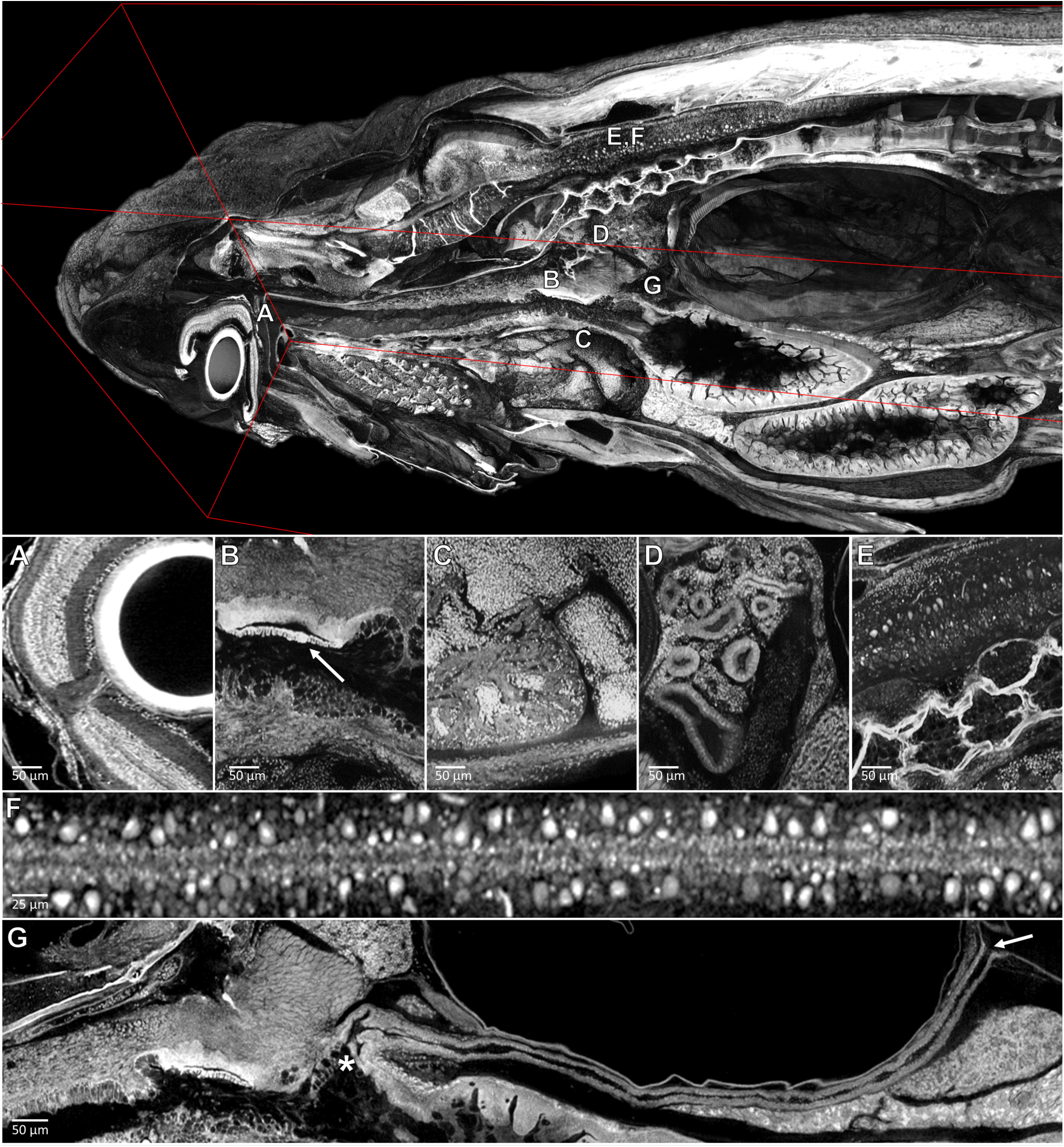
Pan-cellular staining and variable-thickness views allow for characterization of 3D microanatomical features. *Top:* Cutout visualization of a juvenile (33 dpf) zebrafish stained with PTA showing detail in many soft tissue structures. *Bottom*: Cell types and structures that can be visualized include neuronal cells in the eye (A), cartilaginous rudiments of the squamous patch dorsum (arrow) of the pharynx (B), nucleated red blood cells and heart chambers (C), nephrons of the kidney (D), brain nuclei and motor neurons in spinal nerve cord (E, F) and the length of the pneumatic duct (proximal and distal ends noted by an “*” and arrow, respectively) (G). Panels A and E represent individual slices (1.43 μm in thickness) while panels B, C, D represent maximum intensity projections of 5 μm thick sections to visualize larger 3D structures. F represents a 7 μm thick projection.

### Detection of Histopathological Features

Successful histopathological analysis requires the detection of subtle differences in staining at high resolution correlated with a range of changes in specific micron-scale structures. To probe the ability of our approach to detect histopathological change, we used a mutant, *huli hutu* (*hht*), that shows histologically-visible morphological change across all cell types and tissues (*30*) (**Figure S5**, **Movie S6**). Reconstructions of larval *hht* at 0.743 µm^3^ voxel resolution allowed us to detect all of *hht*’s known histological changes, including cell death (nuclear fragments associated with karyorrhexis), nuclear atypia (increased size, deviation from typical ovoid shape, and irregular nuclear membrane contour) in the gut, and tissue degeneration. Moreover, we were able to establish the absence of the pneumatic duct, by inspecting every virtual slice of the entire fish at sub-micron resolution. This structure cannot be reliably studied in histology on account of its small size and the variability in fish alignment prior to sectioning. Age matched wild-type and *hht* fish (2, 3, 4 and 5 dpf) are available on our web-based data sharing interface to qualitatively track progression of histopathological change (**Table S1**).

### ViewTool Enables Web-based Access and Data Sharing

The file sizes associated with synchrotron micro-CT are on the order of ∼100 GB per larval or juvenile zebrafish, making scans difficult to view for people with standard computational resources. To allow users to inspect the data without having to download the full resolution volumes, we developed ViewTool, an open-access and web-based multiplanar viewer (http://3D.fish) (**Figure S6**). ViewTool allows for accurate visualization of micro-anatomic features of normal and abnormal tissue in three orthogonal planes. Coronal and sagittal image stacks were digitally resliced from the original 2D axial reconstructions. While re-slicing of the 3D file at any angle is possible, ViewTool allows the user to explore a full volume through synced interrogation of three orthogonal projections. Any single slice can be examined at full-resolution in a pop-up zoom interface. ViewTool combines both radiology and digital pathology workflows into a seamless experience while offering a low barrier of entry to our 3D data.

### Cell Nuclei Distribution Reveals Phenotypic Variation in Larval Zebrafish Brain

Due to the combination of our field-of-view and resolution, we are able to compute features of individual cells and pattern of cells throughout the whole specimen. Anatomic pathologists use these cytological characteristics in qualitatively tissue assessments (*31, 32*). Many of these attributes can be quantified in 3D, which would bring added value in distinguishing disease states and understanding normal tissue and cellular architecture. Indeed, the detection of phenotypic abnormality requires knowledge of the spectrum of normal samples. To evaluate the resolving power of histo-tomography to detect individual phenotypic variation, we combined regional segmentation with automated cell detection for characterization of the zebrafish brain (**Figure S7**).

We conducted shape (elongation) and size (volume) measurements of manually segmented (n=20) nuclei in 5 dpf zebrafish (**Figure 6A-B**). As anticipated, measurements of shape and size of red blood cells (RBCs) and motor neurons were different, in agreement with the clear morphological differences apparent in histology (**Figure 6C**). In addition, RBCs and motor neurons are more elongated versus typical brain nuclei, which are more spherical. Specifically, RBCs are known to have a flat and oval morphology while motor neurons are more “tear drop” shaped. Other differences include the volume, with motor neurons having the largest average size and typical brain nuclei the smallest. Note that the cytoplasm of RBCs and motor neurons stains darkly with PTA, unlike that of other brain cell nuclei. From these measurements, it is apparent that our current resolution is sufficient in distinguishing brain nuclei from larger classes of stained objects.

**Figure 6.**
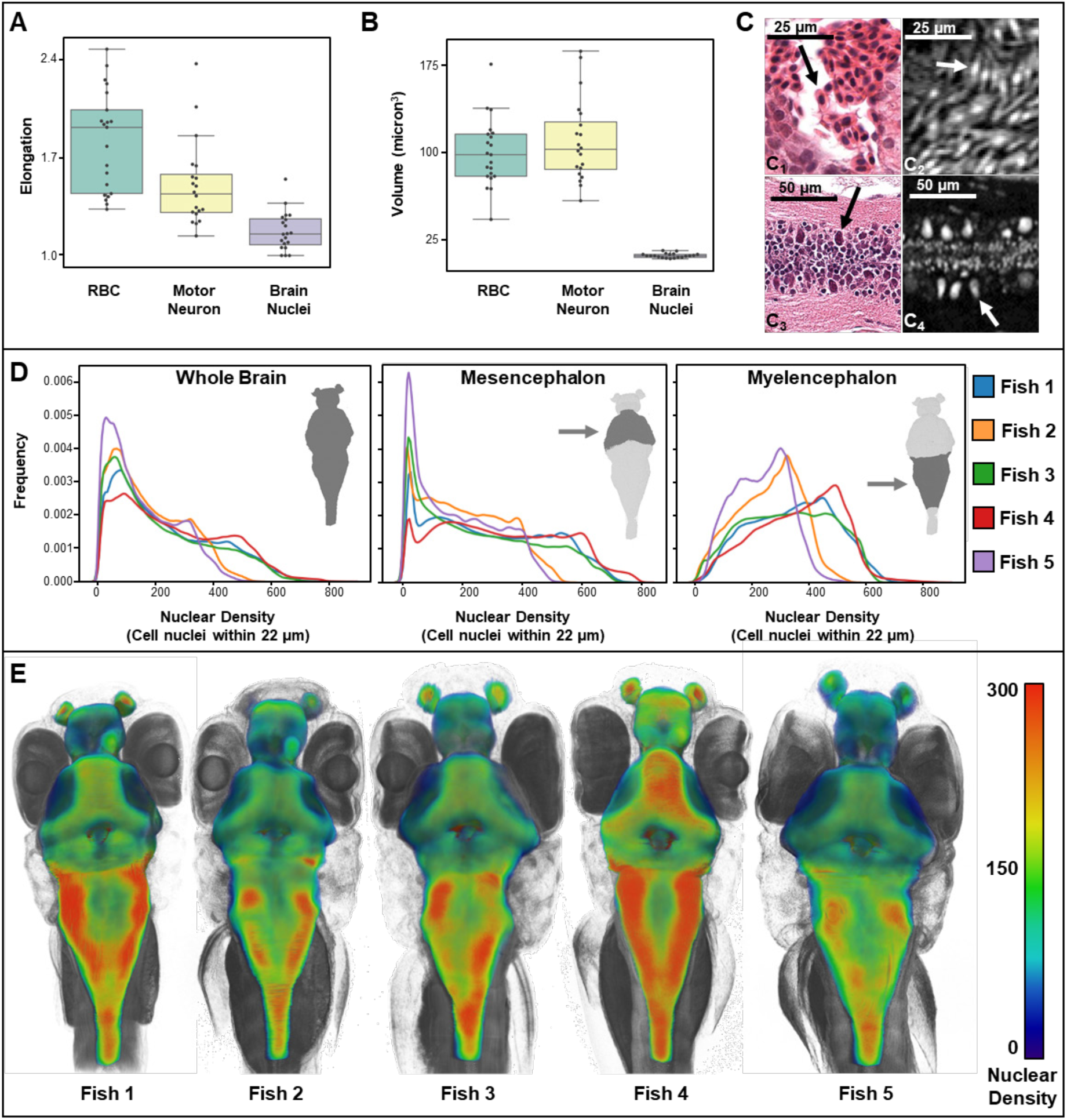
Cell nuclei distribution in larval zebrafish reveals phenotypic variation. Cytology (i.e., elongation and volume) of manually detected brain cell nuclei varies from that of red blood cells (RBCs) and motor neurons (n=20 cells per condition) (A, B). Differences are consistent with what is visually apparent in both histology (C1 and C3) and histo-tomography (C2 and C4) for RBCs and motor neurons, respectively. Differences in cell nuclei distribution can also be observed. Cellular density varies between individual brain regions and carries distinct signals consistent between individual samples (D). A 3D rendering of whole-brain densities is presented for each fish (E) and reflects signal differences presented in (D).

We have been able to assign brain nuclei to major brain regions including the olfactory epithelia, telencephalon, diencephalon, hypothalamus, mesencephalon, metencephalon, myelencephalon, white matter, and spinal cord (*33*–*35*). Cell nuclei were automatically detected from head scans of intact fish using a supervised learning approach with a random forest classifier. Three 75 µm^3^ regions of neural cell nuclei were manually annotated and compared against classifier results for location and counts. The F_1_ score, optimized for probability threshold to balance precision and recall, showed ∼90% correspondence between manual and automated detections (**Figure S8**). The patterns of cell nuclei in our samples corresponded well with those seen in 54 nm thick transmission electron microscopy sections (*9*) (**Figure S9**).

Quantitative analyses of cytological features in the brain demonstrate robust individual and collective cell measures. The average number of nuclei in the brain region studied was 75,413 (SD = 8,547, n=5 fish) (**Table S2**). The size and shape of populations of cell nuclei were compared between individuals. Nuclear size and shape showed no significant differences in whole brains between five replicate specimens (**Figure S10**). In contrast, we observed differences in nuclear density (number of detected cell nuclei within 22 µm of an individual voxel) between brain regions, where individual density distributions and 3D patterning showed variation between wild-type specimens (**Figure 6D-E**). Comparatively, fish 2 and 5 exhibit a left shifted density distribution versus fish 1, 3, and 4, in association with lower cell counts and larger volumes in fish 2 and 5 (**Tables S2-3**). In addition, prominent differences in cell density were seen in the olfactory epithelium, metencephalon and myelencephalon, possibly associated with slight differences in exact developmental stage.

## Discussion

Large-scale studies of the effects of genes and environment on phenotype (so-called phenome projects) are ideally comprehensive and quantitative in nature, covering all cell and tissue types. Given the desire for whole-organism, complete histological phenotyping, we pursued synchrotron micro-CT as a way to maximize the ability to phenotype small organisms in 3D, at a throughput and resolution that is compatible with phenome projects (*36*). High-throughput adaptations of histo-tomography in zebrafish will make it possible to quantify morphological changes caused by mutations in each protein-encoding gene (>26,000) (*37*). Over 70% of human protein-coding genes have at least one zebrafish orthologue. Therefore, human gene function can be probed by studying the morphological phenotypic consequences of single gene defects in zebrafish. Such an effort could be the basis of a Zebrafish Phenome Project with purposes comparable to those of established mouse phenome projects (*38, 39*). Our synchrotron micro-CT pipeline may be particularly well-suited for phenome projects due to sample stability (*26*) and acquisition throughput. By our estimates, screening 20,000 mutants in replicate would require ∼40 years with monochromatic source and ∼1 year using pink-beam. A high-throughput primary screen by pink-beam could be followed by higher-quality monochromatic imaging of samples showing evidence of phenotypic change.

An ideal of comprehensive histological phenotyping would include quantitative characterization of tissue architectural and cellular features beyond simple measures such as organ size and body shape. The combination of resolution, field-of-view, pan-cellular staining, and throughput of our approach enables this ideal across multiple whole samples. Importantly, the ability to compute collective features such as cellular density has allowed for detection of individual phenotypic variation, as illustrated in the brain. The addition of tissue-, cell-, and protein-specific micro-CT stains will enable targeted analyses and cross-atlas comparisons of explicit constituents within the full context of a scanned sample or organism.

The work shown here approaches the ideal of comprehensive and quantitative phenotyping for model organisms that lie in the mm-to-cm length scale such as the zebrafish. Indeed, synchrotron X-ray micro-CT allows for full volume imaging of replicate samples at cellular resolution, distinguishing neighboring cells and resolving nuclear morphologies in optically opaque juvenile zebrafish specimens. Extending this work to *Drosophila, C. elegans*, and *Arabidopsis*, and to tissue samples from larger organisms such as mouse and human would enable analyses across species and have the potential to lead to a more universal understanding of variations in tissue architecture and morphological phenotype. To make these data accessible to users with standard computational resources, we have developed an open-access, web-based data sharing platform, ViewTool, that allows both 2D and 3D histological analyses of full sample volumes. Repositories of high-resolution histo-tomographic images from a full complement of human and other organismal tissues would facilitate cross-correlations between model system and human phenotypes (*40, 41*). Such a resource would have the potential to increase analytical precision, sensitivity, reproducibility, and data sharing to address important questions across basic and clinical sciences.

## Acknowledgments

The authors would like to thank Steven Peckins and Drs. Timothy Cooper and Gordon Kindlemann for critical insight in the early stages of the project’s conception and development, and Will Fan for proofreading. **Funding:** The investigators acknowledge NIH funding support (PI: KCC, R24-RR017441, and PI: KCC, R24-OD018559) and pilot award funding to KCC from the Huck Institutes of the Life Sciences and the Institute for Cyber Science, PSU. The use of the APS, an Office of Science User Facility operated for the US Department of Energy (DOE) Office of Science by Argonne National Laboratory, was supported by the US DOE under contract no. DE-AC02-06CH11357. **Author Contributions:** KCC envisioned and defined the goals of the project in consultation with PLR, who proposed and enabled the use of Argonne National Laboratory’s micro-CT instrumentation at 2-BM. X Xiao and FDC at Argonne National Laboratories, and YW, defined the basic parameters of synchrotron micro-CT with PLR. KCC defined specimens, sample preparation, and parameters for imaging results. KCC, DC, YD, X Xiao, and FDC conducted imaging experiments and analyzed initial results with DC and XX. KCC, AYN, JEC, and KCA oversaw zebrafish husbandry and sample preparation. YD, DJV, MAY, and PV conducted all computational experiments, and interpreted results in consultation with KCC, PLR, YW, VAC, KCA, and DBvR. YD, DJV, MAY, and SRK created figures in consultation with KCC, KCA, VAC, and DBvR. DJV and XXin created the movies motivated by and in consultation with KCC. KCC initiated and DJV created ViewTool in consultation with KCC, YD, MAY, SRK and DBvR. YD, DJV, MAY, DBvR, SRK and KCC wrote the manuscript, with input and edits by all authors. **Competing Interests:** All of the authors declare that there are no conflicts of interest. **Data Availability:** Zebrafish reconstructions are available through our online open-access multiplanar viewer platform, ViewTool (http://3D.fish). Full resolution datasets, including raw projection data, are available by request as a download or physical media.

## Supplementary Materials List

Materials and Methods

Figures S1-S10

Tables S1-S3

Movie Captions S1-S6 Supplemental References (#41 - 51)

## Abbreviations

3D: three-dimensional
H&E: hematoxylin & eosin
2D: two-dimensional
micro-CT: X-ray micro-tomography
dpf: days post-fertilization
PTA: phosphotungstic acid
RBC: red blood cell

